# Stable germline transgenesis using the *Minos* Tc1/*mariner* element in the sea urchin, *Lytechinus pictus*

**DOI:** 10.1101/2024.03.26.586777

**Authors:** Elliot W. Jackson, Emilio Romero, Svenja Kling, Yoon Lee, Evan Tjeerdema, Amro Hamdoun

## Abstract

Stable transgenesis is a transformative tool in model organism biology. While the sea urchin is one of the oldest animal models in cell and developmental biology, it has relied on transient manipulations of wild animals, and has lacked a strategy for stable transgenesis. Here we build on recent progress to develop a more genetically tractable sea urchin species, *Lytechinus pictus*, to establish a robust transgene integration method. Three commonly used transposons (*Minos, Tol2, piggyBac*) were tested for non-autonomous transposition, using plasmids containing a polyubiquitin promoter upstream of a H2B-mCerulean nuclear marker. *Minos* was the only transposable element that resulted in significant expression past metamorphosis. F_0_ animals were raised to sexual maturity and spawned to determine germline integration, transgene inheritance frequency, and to characterize expression patterns of the transgene in F_1_ progeny. The results demonstrated transgene transmission through the germline, the first example of a germline transgenic sea urchin, and indeed of any echinoderm. This milestone paves the way for the generation of diverse transgenic resources that will dramatically enhance the utility, reproducibility, and efficiency of sea urchin research.

**Significance Statement:** Transgenic tools are essential for effective utilization of animal models. Despite being an established model for cell and developmental biology, the sea urchin has not previously benefited from transgenic technology. This study reports the generation of the first germline transgenic sea urchin and opens new avenues for this organism.

## Introduction

Sea urchins are classical models for cell and developmental biology. Sea urchins sit at an informative phylogenetic position as basal deuterostomes and offer specific biological advantages as animal models, including extreme fecundity, optical transparency, and clockwork synchrony of development. Research in these animals has informed mechanisms of fertilization, cell division and early development. Examples include the discovery of cyclins (1), which ultimately led to currently approved drugs targeting metastatic cancers (2), and the first descriptions of gene regulatory networks (3, 4), which have become a foundational framework for understanding development (5–7). Sea urchins remain favorites for study across diverse fields including gene regulation of development (8–10), innate immunity (11–16), biomineralization (17–21), nervous system development (22–24), reproduction (25–27), and developmental physiology (28–30).

Despite the history and importance of sea urchins, they have been bottlenecked by the lack of a coherent strategy for development of stable genetic lines. This has resulted in the lack of germline transgenics. Prior approaches for expression of transgenes in sea urchins have relied upon transient manipulations of wild animals. These methods are labor intensive, less reproducible and are limited in utility to the early stages of development. The status quo for expressing transgenes is microinjection of reagents such as synthetic mRNAs or bacterial artificial chromosomes (BACs). Each experiment typically uses embryos from a new male/female wild-type mate pair and is terminated well before an F_1_ generation is reached. This is in stark contrast to the renewable nature stable genetics, defined as germline modifications in a F_1_ population and beyond, which are the gold-standard in forward and reverse genetic approaches.

As with several other model organisms, there are multiple species of sea urchins used by researchers. Most were chosen long ago, based on local availability of wild animals. Many have long generation times and/or specialized culturing requirements making them poorly suited for culturing in the lab. A notable exception is *Lytechinus pictus*, which is native to the California coast, has a long history in the field, and is suitable for egg-to-egg culture in the lab (31). We have previously shown that *L. pictus* has a generation time of 4-6 months, grows at room temperature (∼18-22°C), and has a modest adult size suitable for culturing in standard recirculating systems (32). As such, *L. pictus* is one of only two species worldwide which have been used to produce homozygous mutant lines using CRISPR/Cas9, a key step towards stable genetics (33, 34). This has left transgenesis as the final barrier towards a genetically enabled sea urchin.

In this study, we report the generation of the first germline transgenic sea urchin using transposon-mediated integration in *L. pictus*. To achieve this, a polyubiquitin promoter driving expression of a fluorescent nuclear marker was used to test three DNA transposons for non-autonomous integration. We identified *Minos* Tc1/*mariner* as the only transposable element that resulted in significantly higher integration rates as compared to controls. We then leveraged this finding to compare the transgene expression patterns across larval development to determine differences between transient expression of the plasmid and stable transgene expression from the genome. Finally, we examined transgene expression patterns between F_0_ and F_1_ progeny. This study paves the way for a new era in the generation of diverse sea urchin lines useful in cell fate analysis, live imaging, conditional gene expression, and combinatorial mutagenesis.

## Results

To design a suitable transgenesis strategy for sea urchins, we first needed to discover a reporter with strong ubiquitous expression for efficient screening of transformants. Using prior expression data, we selected putative promoters that express strongly in both larval and juvenile phases, to ensure ability to readily select juveniles with somatic integrations. Of the five promoters tested, the promoter of an *L*.*variegatus* polyubquitin-C gene (LOC121415894), cloned upstream of a cyan fluorescent protein (CFP) mCerulean (LvPolyUb::H2B-CFP) was the only one that expressed reliably. This promoter element is 3,439 bp and spans approximately - 1000 nt upstream of the transcriptional start site to the start codon in the *L*.*variegatus* genome (**Figure 1**).

**Figure 1.**
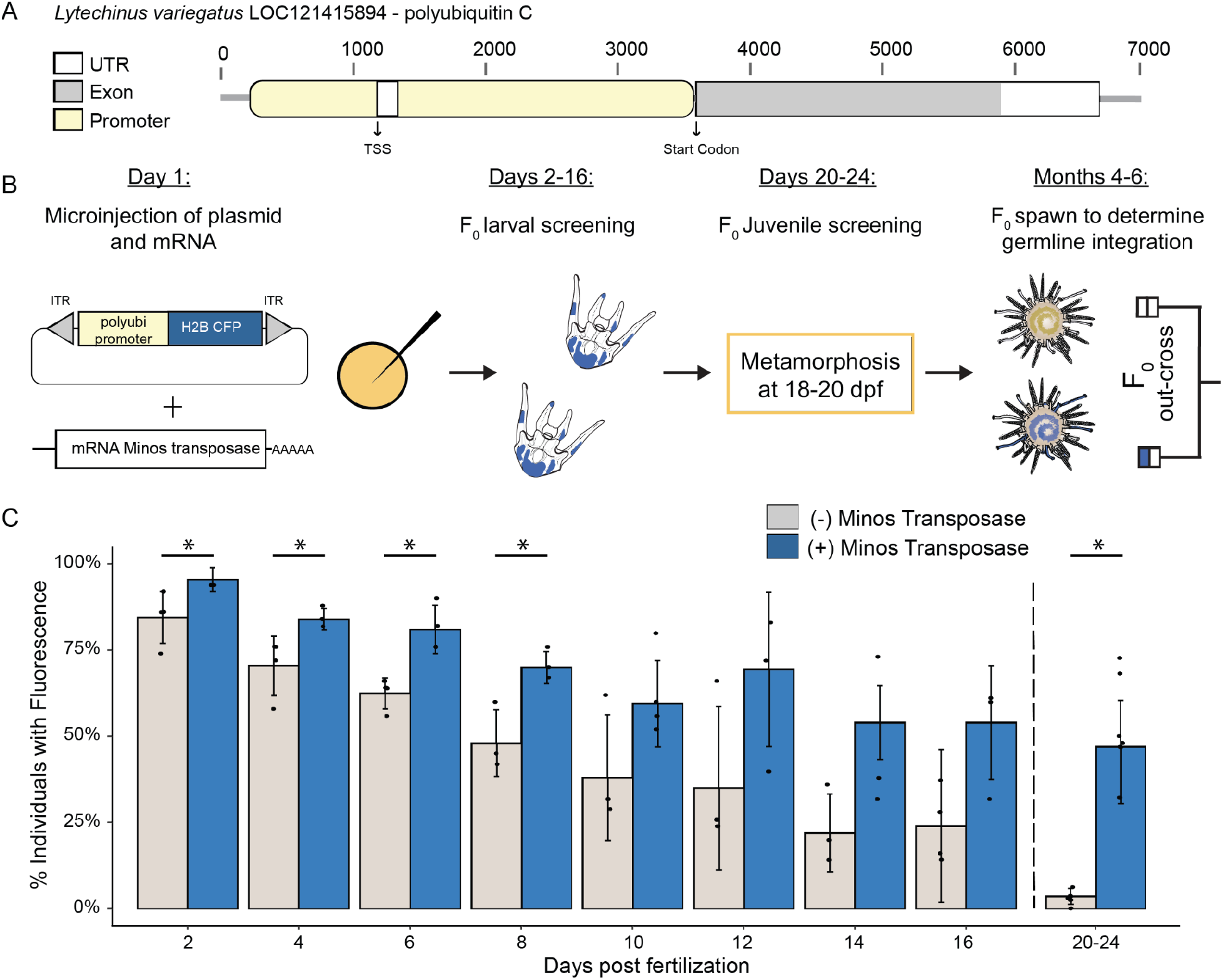
Integrated transgenes are expressed robustly at the late larval and juvenile stages. Transient transgene expression from the reporter construct alone was distinct from genomic transgene expression from *bona fide* integration at the late larval and juvenile phases.(A) Schematic of the polyubiquitin promoter from *L*.*variegatus* used to drive transgene expression. (B) Experimental design for creating F_0_ sea urchins with somatic integrations that could subsequently be selected for grow out. (C) Transgene expression patterns with and without the *Minos* transposase over early development. Each data point represents the proportion of offspring with fluorescence from a unique mate pair. In total 200 larvae were screened for each timepoint. Error bars represent one standard deviation from the mean. Black dotted line represent the transition from larval to juvenile through metamorphosis. Asterisks indicate significant differences using Welch two-sample *t* test (*P* <0.05).

Using this construct with the *Minos* transposase, which had previously been demonstrated to achieve effective transgenesis in a related invertebrate, *Ciona intestinalis* (35), we took advantage of the rapid development of *L. pictu*s to screen for integration. In these experiments we compared the change in fluorescence across larval development and shortly after metamorphosis between embryos injected with only the reporter construct to embryos injected with the reporter construct and mRNA of the *Minos* transposase **(Figure 1**). It has been previously shown that exogenous DNA persists through larval development (36) and is largely cleared after metamorphosis. Based on this, we expected differences between transient forms of expression (driven by the plasmid itself) and those driven by *bona fide* integrations into the genome, would be most pronounced after metamorphosis.

In control larvae injected with just the reporter construct, observable fluorescence declined by ∼10% every two days until reaching 3.6% in animals post metamorphosis. In larvae with both the reporter construct and *Minos* transposase, the percentage of larvae with fluorescence also dropped approximately 10% every two days until 14 days post fertilization stabilizing around 50% (**Figure 1**). Consistent with our expectation, the most significant change in the percent of individuals with fluorescence between experimental (47.4%) and control (3.57%) animals was after metamorphosis, 43.83% difference (Welch two-sample *t* test, *P* = < 0.0001, *df* = 5.28, *t* = 8.07). Statistically significant differences between experimental groups were also found from days 2 through 8 (Welch two-sample *t* test, *P* = < 0.05), though the observed qualitative differences between groups (i.e. number of cells expressing the transgene and brightness) was difficult to distinguish until close to metamorphosis (**Figure 1**).

Next we determined whether other transposases might also be effective at introducing similar, stable somatic integrations. *Tol2* and *piggyBac*, which are two commonly used transposable elements used to generate transgenic animals, were tested. We generated LvPolyUb::H2B-CFP constructs for each of these and tested their effectiveness at introducing somatic integrations in juveniles. Neither *Tol2* (3.0%, n=714) or *piggyBac* (1.9%, n=426) yielded integration rates significantly different than control conditions (3.6%, n=532) (Pairwise Wilcoxon test, *P* = > 0.05) (**Figure 2**). *Minos* was the only transposase with significantly different somatic integration rates compared to control conditions, *Tol2*, and *piggyBac* (Pairwise Wilcoxon test, *P* = < 0.05). The efficiency of somatic integration using *Minos* ranged from 32-72% with an average of 47% (n=883).

**Figure 2.**
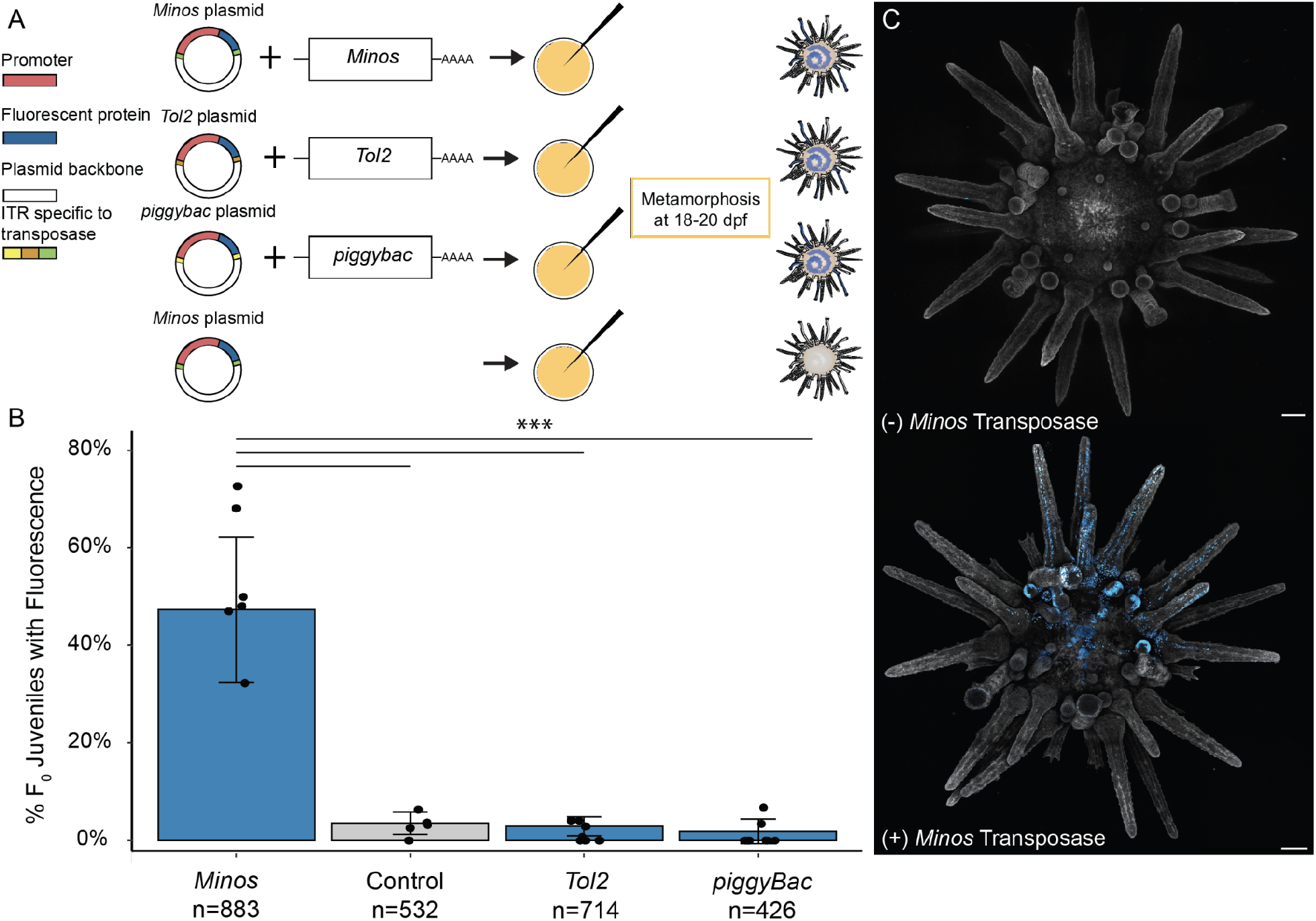
The *Minos* element resulted in significantly higher somatic integration rates than *Tol2, piggyBac* or controls. *Tol2* and *piggyBac* were not significantly different than controls. (A) Each transposon was tested by injecting mRNA of the transposase along with a plasmid containing a polyubiquitin promoter upstream of a human H2B-mCerulean. Control conditions only contained the plasmid. (B) Screening for somatic integration was performed 2-4 days post-metamorphosis. Each data point represents the proportion of offspring with fluorescence from a unique mate pair with error bars showing one standard deviation from the mean. Asterisks indicate significant differences between groups (Pairwise Wilcoxon rank sum test, Bonferroni-corrected *P* < 0.0001). (C) F_0_ juveniles injected with the reporter plasmid and with (+) or without (-) the *Minos* transposase. Juveniles were stained with CellMask plasma membrane orange to create contrast with nuclear CFP signal. Scale bar = 100μm.

Given these results we used *Minos* to produce transgenic adults to assay for germline integration. F_0_ juveniles exhibited strong, mosaic transgene expression patterns that persisted through to adulthood (**Figure 3**). To determine if the somatic integrations we observed in juveniles might also pass through the germline we raised 70 fluorescent F_0_ juveniles through to sexual maturation. At approximately 4 months after metamorphosis, we qualitatively graded these individuals based on their fluorescence. Of these, 48 had moderate to strong, mosaic transgene expression patterns. These 48 animals were subsequently spawned and crossed with wild type animals to determine if their gametes produced transgenic larvae. Of the 48 individuals, we obtained 27 males and 21 females. Interestingly, we observed that sperm from 10 of the males produced transgenic larvae whereas eggs from only two females produced transgenic larvae. To determine whether this might be mediated by some form of female-specific transgene silencing we tested for the presence of integration by PCR. We found that all animals that produced progeny with observable transgene expression were also positive by PCR, and all animals without observable transgene expression were negative by PCR (**Supplemental Figure 1**).

**Figure 3.**
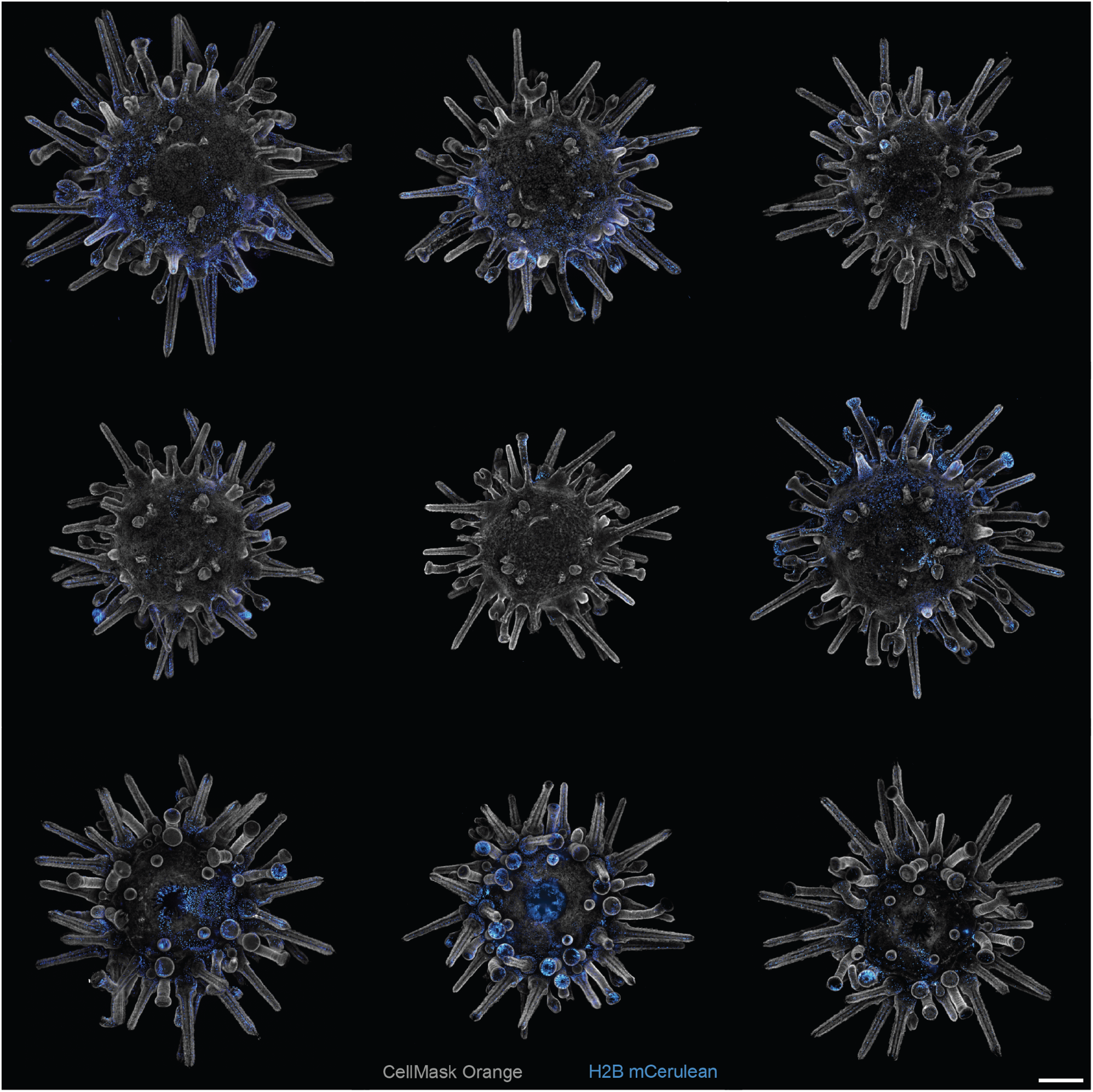
Mosaicism and variation of transgene expression in F_0_ animals. Live imaging of nine unique F_0_ juveniles injected with *Minos* transposase and our reporter construct. Top and middle row is the aboral view and the bottom row is the oral view. Juveniles were stained with CellMask plasma membrane orange to create contrast with nuclear CFP signal. Scale bar = 250μm.

We then sought to determine the level of transgene expression across individuals with germline integrations, determine the timing of transgene expression and characterize transgene expression patterns in F_1_ larvae and juveniles. To determine the percentage of transgenic offspring produced from each individual with germline integration we used a high-content imaging approach (37). We screened 353-1295 F_1_ embryos from each outcross of the 12 F_0_ animals with germline integration (**Figure 4**). The average germline transmission rate was 41.81% (n=3826/8579) and varied from 6.26% to 64.30% (**Table 1**).

**Table 1.** Proportions of F_1_ progeny expressing integrated transgenes. F_0_ transgenic animals with germline integrations were outcrossed to wild-types in order to determine the proportion of offspring expressing the transgene.

| **Animal ID** | **Integrated** | **Non-integrated** | **Total Screened** | **% integrated** |
| --- | --- | --- | --- | --- |
| #01 Male | 311 | 680 | 991 | 31.38% |
| #13 Male | 39 | 299 | 338 | 11.54% |
| #15 Male | 249 | 658 | 907 | 27.45% |
| #21 Male | 697 | 598 | 1295 | 53.82% |
| #22 Male | 523 | 449 | 972 | 53.81% |
| #28 Male | 518 | 387 | 905 | 57.24% |
| #30 Male | 415 | 342 | 757 | 54.82% |
| #36 Male | 225 | 142 | 367 | 61.31% |
| #43 Male | 97 | 256 | 353 | 27.48% |
| #46 Male | 381 | 347 | 728 | 52.34% |
| #09 Female | 27 | 404 | 431 | 6.26% |
| #52 Female | 344 | 191 | 535 | 64.30% |

**Figure 4.**
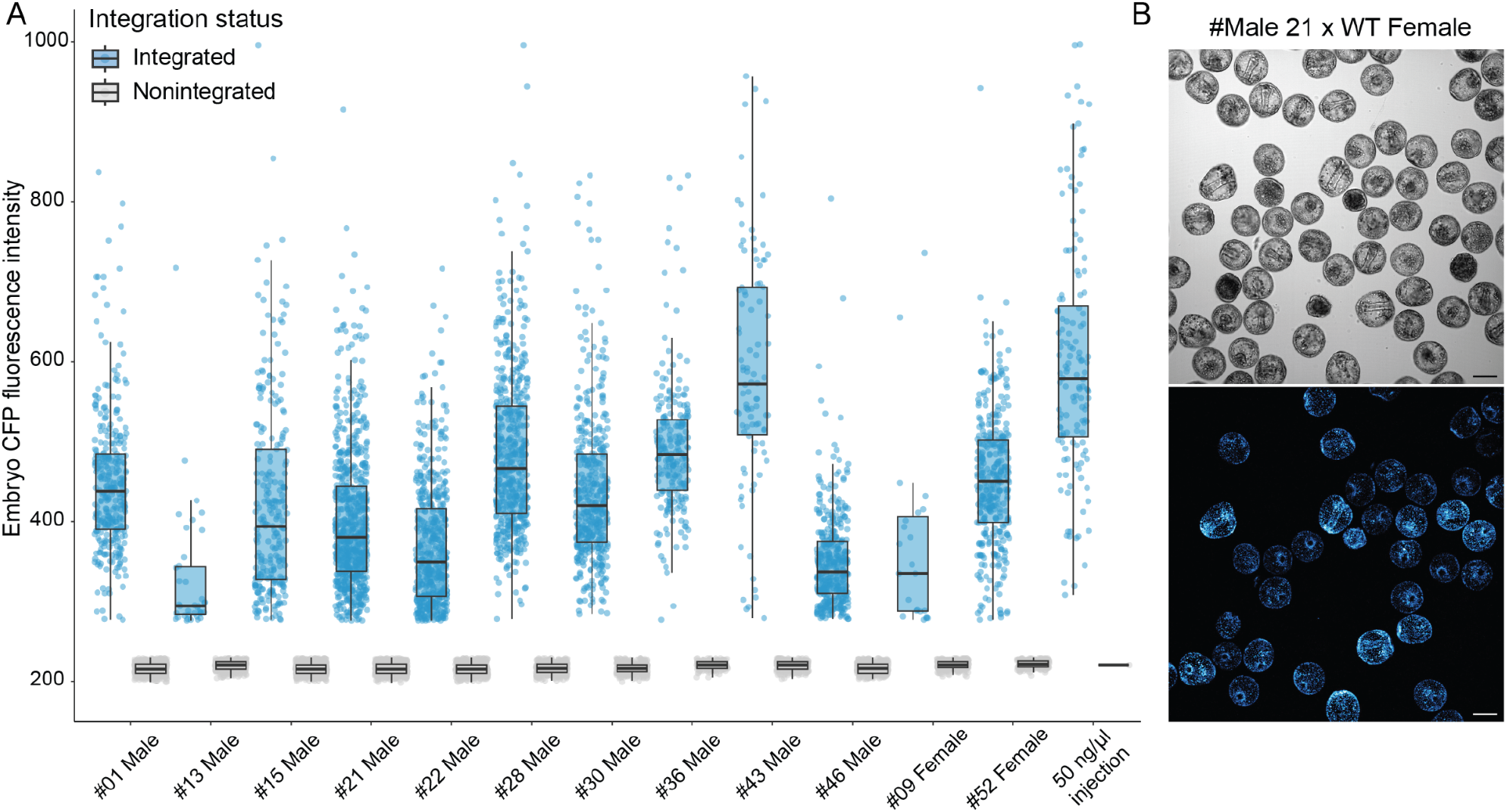
Measurement of transgene expression levels by high content screening of F_1_ embryos. (A) Max fluorescent intensity of transgenic embryos separated by parent with germline integration. Each dot represents one embryo 24-26 hours post fertilization. (B) Representative image used for machine learning-based automated segmentation to determine integration status and measure CFP fluorescent intensity. The top and bottom images are the same embryos in transmitted light and confocal, respectively.

Next, we examined the relative level of transgene expression across individuals and compared transgene expression to overexpression of histone CFP by injection of mRNA, which is the current standard methodology for expression of a histone marker in sea urchins (**Figure 4**). All transgenics produced brightly fluorescent embryos that could be easily distinguished from controls by eye, although transgene expression varied 2.5-fold from the brightest to the dimmest individuals. Of these the brightest transgenic embryos were in the range of a 50 ng/μl mRNA overexpression concentration (**Figure 4**) and were more than adequate for visualization in routine confocal microscopy.

Finally, we examined the timing at which the transgene becomes expressed and whether it persists through metamorphosis in F_1_ progeny. Whether driven through the male or female germline, expression of this construct in F_1_ progeny begins at hatching (∼8 hpf at 22°C) and persists through development. Unlike transient overexpression, nuclei remained brightly labeled through larval development including in competent larvae that were approaching settlement (**Figure 5**). Similarly, transgene expression persisted after settlement and into juveniles of the F_1_ generation with variation among individuals in relative intensity (**Figure 6**).

**Figure 5.**
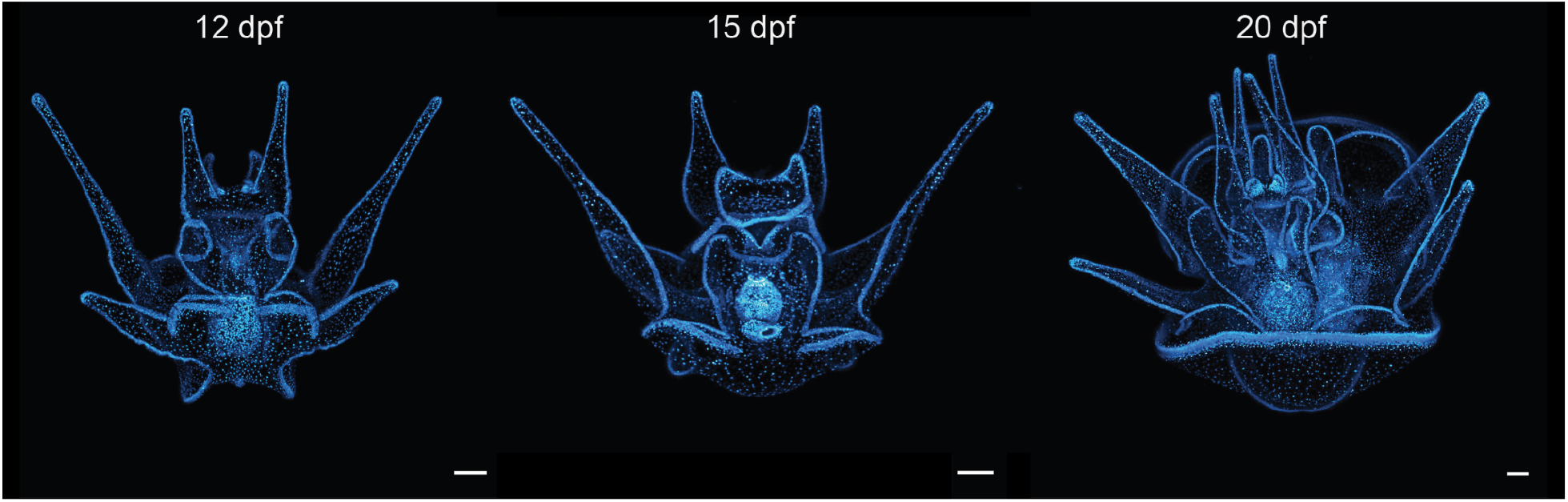
Transgenes are robustly and ubiquitously expressed in F_1_ larvae through metamorphic competency. Image shows confocal micrographs of F_1_ larvae produced from mating wild-type female eggs with sperm of a transgenic male with germline integration. Scale bars = 100μm.

**Figure 6.**
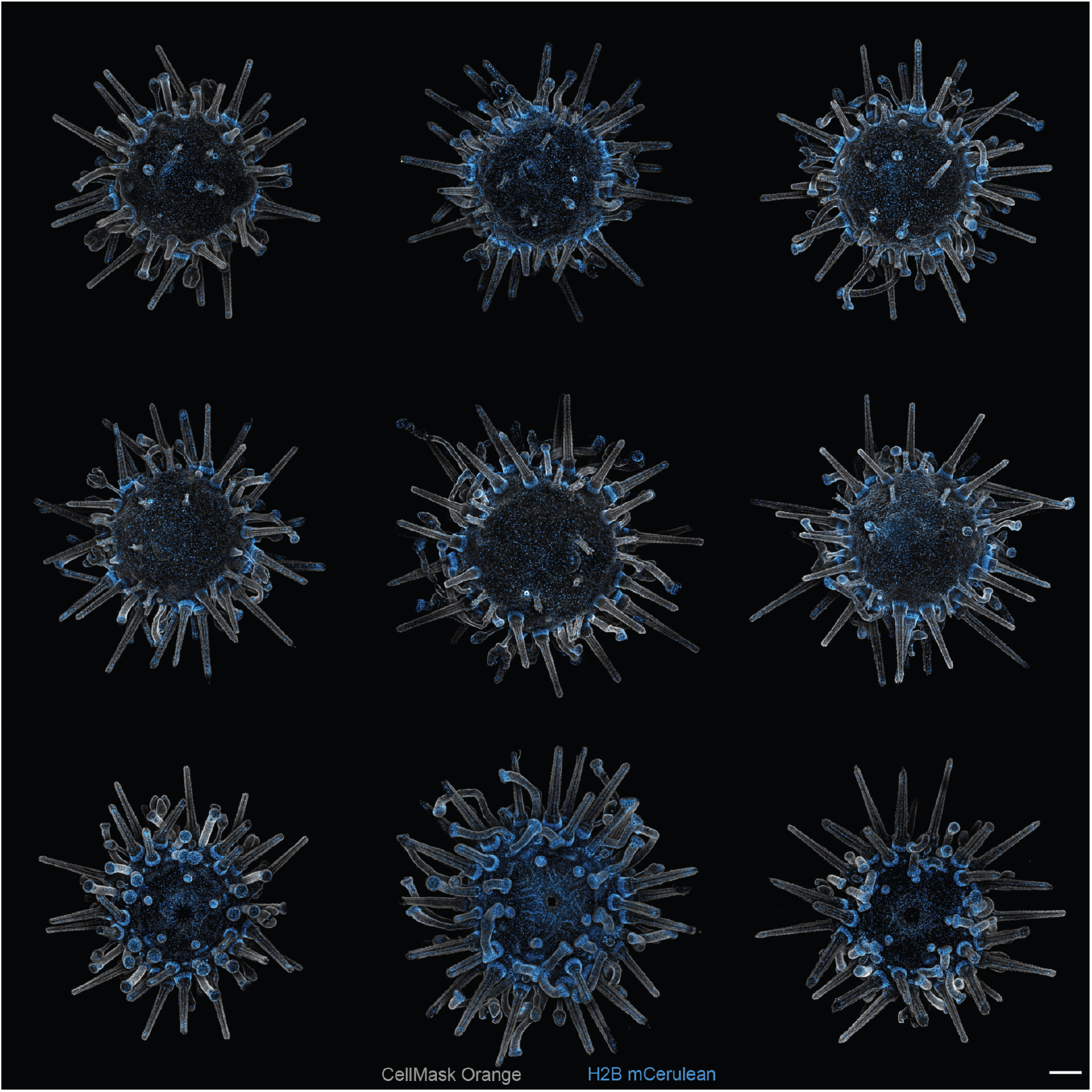
Transgene expression in F_1_ animals is ubiquitous across the body. Live imaging of nine unique F_1_ juveniles. Top and middle row is the aboral view and the bottom row is the oral view. Juveniles were stained with CellMask plasma membrane orange to create contrast with nuclear CFP signal. Scale bar = 250μm.

## Discussion

This is the first study to demonstrate stable germline transgenesis in sea urchins, or indeed in any echinoderm. Prior to this study there was no consensus about how to stably integrate exogenous DNA into the genome of a sea urchin. The first attempt at transgenic manipulation in sea urchins was performed by microinjection of naked DNA (36, 38). Both circular and linearized plasmid DNA were microinjected into embryos with only a few percent of the injected sea urchins retaining the injected constructs past metamorphosis, and germline integration was not demonstrated. We similarly observed very low levels of fluorescence in juveniles injected with plasmids in the absence of transposases (**Figure 2**). However, when using the *Minos* Tc1/*mariner* element we observed an order of magnitude greater rate of integration then injecting a plasmid alone (**Figure 2**). This reliably produced high levels of somatic integration in roughly 50% of the injected F_0_ juveniles. Somewhat unexpectedly, somatic integration rates using *Tol2* and *piggyBac* were nearly the same as injecting the plasmid alone, suggesting that these transposons are not effective in sea urchins.

One of the biggest obstacles for developing a transposon-based integration method in sea urchins is being able to delineate between expression from the injected plasmid versus expression driven from a stably integrated transgene. This is because sea urchins, and other echinoderms, will readily express genes on exogenous DNA, whether injected as circular or linear DNA (39). Our results indicate that around 12 days post fertilization in *L. pictus*, the difference becomes more apparent (**Figure 1**). Two qualitative differences between *Minos* injected larvae and controls are the level of fluorescence and number of fluorescent cells, both of which appear higher in larvae injected with the *Minos* transposase. We assume that the most likely reason for this is the dilution of the circular plasmid with cellular division, leading to a weaker fluorescent signal as the animal grows (36). However, we found screening for integration was ultimately most conclusive in juveniles because transient plasmid expression nearly disappears at metamorphosis (**Figure 2**).

In this study we noted some possible differences in germline transmission between males and females. The most obvious of these was the disproportionate number of males compared to females with germline integration. The second observation, which may or may not be related to the first, is the difference of transgene variegation of expression (**Supplemental Figure 2**). Specifically, we observed more variegation in the fluorescence levels of F_1_ larvae from females. We speculated that this might be due to some sort of female specific silencing of the integrated transgene in females. However, our analyses did not support this hypothesis, as all females lacking the transgene expression, also lacked integration by PCR. Thus, alternative explanations might be that the female germline is somehow more resistant to integration, or that this difference was simply an artifact.

In both males and females transgene expression was observed upon hatching (∼8 hpf at 22°C), which is when the major wave of zygotic gene expression occurs in sea urchins (27, 40). This was unexpected as the publicly available RNA-seq data which show polyubiquitin transcripts present in the egg (**Supplemental Figure 3**) and recruited to polysomes at fertilization (27). One possible explanation for this is the polyubiquitin-C 3’UTR element, which was not a part of the transgene cassette that may include regulatory motifs responsible for maternal deposition as well as stability and translation during early embryogenesis (41). However, for other ubiquitously expressed genes such as ubquitin, maternal deposition of eGFP into zebrafish eggs was ultimately observed in the F3 generation using a promoter excluding the 3’UTR element (42). Thus, it is conceivable that egg expression of our transgene might be observed in subsequent generations of this line.

Stable transgenesis has the potential to reshape the landscape of sea urchin research. Stable transgenesis offers significant advantages over less-reproducible and repetitive transient methods currently used in the field. For example, this study demonstrates how marker transgenes, such as histone or membrane markers, can be introduced by simply fertilizing wild eggs with transgenic sperm rather than by overexpression of mRNA (43, 44). Since sea urchins make abundant gametes, and *L. pictus* sperm can easily be cryopreserved or extended (45) virtually any lab will be able to easily make a surplus of transgenic embryos by simply fertilizing eggs with transgenic sperm. In addition, we have shown that the marker transgenes persist into later development, opening the door to live imaging of later larval and rudiment features that have until now only been accessible by fixation (46). Perhaps the most significant advantage of all, is that transgenesis takes better advantage of the specific biological features of the sea urchin that motivated its use in the first place – namely extreme fecundity and developmental synchrony. Both can be more fully utilized by stable transgenesis where millions of modified embryos can be generated and used in high throughput format (42). This is in stark contrast to the current state of the art where only 10s, or at most 100s of embryos, are generated.

This work is the result of systematic steps to de-risk *L. pictus* as a genetically enabled sea urchin. The outcomes are likely to have significant impact on both the sea urchin and echinoderm community at large, through the production of new animal resources and tools. Importantly this advance opens the door to the generation of diverse sea urchin lines which can be shared among labs and thereby usher in a new era of collaboration. As such, sea urchin transgenics may dramatically impact the throughput, reproducibility and utility of this iconic animal model.

## Materials & Methods

### Lytechinus pictus husbandry

Adult *L. pictus* were collected in San Diego, CA, USA and housed in flowing seawater aquaria at 20-22°C. The materials and methods used to spawn, fertilize, and culture *L*.*pictus* were followed as previously described (34). Juveniles at ∼2 mm size were transferred to a recirculating sea water system (Aquaneering, San Diego, CA, USA) and fed Pacific dulse (*Palmaria palmata*) until they reached ∼1 cm in size at which time they were fed *Macrocystis pyrifera*. F_0_ adults were spawned by injecting 50μl of 0.2 μm filtered 0.55M KCL to collect gametes.

### Promoter selection and testing

Publicly available RNA-seq data on Echinobase (47) from *Strongylocentrotus purpuratus* and *Lytechinus variegatus* were used to find candidate genes for promoter discovery. Candidate genes were chosen based on two criteria, 1) Transcript per million (TPM) values ≥ 200 across all developmental time points and adult tissue types and 2) prior evidence of ubiquitous expression in other animal models. NCBI BLAST was used to identify the homologous *L*.*pictus* genes from *S*.*purp* or *L*.*var* as the query sequence. Based on these criteria, LOC121415894 (polyubiquitin-C) and LOC121418990 (polyA binding protein) from *L. variegatus* were chosen and LOC129259176 (tubulin alpha-1A chain), LOC129261990 (elongation factor 1-alpha), and LOC129265555 (polyA binding protein) from *L. pictus* were chosen for promoter screening. The putative promoter region for each gene was chosen to be between 2000-5000 nucleotides surrounding the transcriptional start site which was synthesized by Twist Bioscience (San Francisco, CA, USA) or GENEWIZ from Azenta Life Sciences (South Plainfield, NJ, USA) and subsequently cloned upstream of a H2B-mCerulean (Addgene #198059). Promoter activity was scored by microinjecting 25ng/μl of circular plasmid and screening for fluorescence in embryos 24 hours post fertilization (hpf). DNA concentrations were determined using a Qubit 4.0 prior to microinjection.

### Transposon plasmid construction, mRNA synthesis, microinjections

Three plasmids were required to make mRNA of each transposase (*Minos, Tol2, piggyBac*). The *Minos* transposase (pBlueSKMimRNA) plasmid was a gift from Michalis Averof (Addgene plasmid #102535). The *piggyBac* transposase (pT7mRNA-PB transposase) was purchased from VectorBuilder (Chicago, IL, USA), and the *Tol2* transposase came from the pCS2FA-transposase plasmid (48). Three separate plasmids were designed and synthesized by Twist Bioscience (San Francisco, CA, USA) or GENEWIZ from Azenta Life Sciences (South Plainfield, NJ, USA) that were compatible with each transposase. These plasmids contained inverted terminal repeats specific to each transposase and a multiple cloning site. Restriction enzyme cloning was used to clone the promoter of an *L*.*variegatus* polyubquitin-C gene (LOC121415894) upstream of a cyan fluorescent protein (CFP) mCerulean (LvPolyUb::H2B-CFP) insert into each transposon plasmid (**Supplemental Figure 3**). All plasmids were sequenced prior to experimentation through services provided by Plasmidsaurus or Primordium Labs.

*In-vitro* transcription using the *Minos* and *piggyBac* transposase plasmids were performed using the mMESSAGE mMACHINE T7 Ultra Transcription Kit (Invitrogen, Waltham, MA, USA) according to the manufacturer’s protocol, and *in-vitro* transcription using the *Tol2* transposase plasmid was performed using the mMESSAGE mMACHINE SP6 kit (Invitrogen, Waltham, MA, USA) following the manufacturer’s protocol. mRNA concentration was determined using a Qubit 4.0 and mRNA quality was determined using an Agilent Tapestation RNA Screentape. Only mRNA with RNA integrity numbers (RIN) 8 or greater was used in experiments. Microinjections were performed on one-cell embryos containing a mixture of 500 ng/μl of transposase mRNA, 20 ng/μl of circular plasmid, and 1ng/μl Rhodamine B dextran. Control injections were performed using the same microinjection solution excluding the transposase mRNA.

### Larval and juvenile screening and for somatic and germline integration

F_0_ screening of embryos, larvae, and juveniles was performed on a Zeiss LSM 700 equipped with Zeiss PI 10 or 20x objectives and a CFP filter with a X-Cite Series 120 Q for fluorescent illumination (Waltham, MA, USA). F_0_ screening of adults prior to spawning were performed on a Leica M165 FC stereo microscope with CFP filter and 10x/23 objectives with a X-Cite Series 120PC Q for fluorescent illumination (Waltham, MA, USA. Animals were subjectively graded (A to D) for strength and area coverage of fluorescence. Animals with strong to moderate (A-C grade) observable fluorescence were subsequently spawned, and outcrossed with a wildtype male or female to determine germline integration. F_1_ embryos were screened 2 hpf (∼4 cell stage) and ∼24 hpf.

A subset of F_1_ embryos from each outcross were also collected ∼24 hpf for DNA extraction and PCR. DNA was extracted using PureLink Genomic DNA Mini kit (Invitrogen, Waltham, MA, USA) following the manufacturers protocol. Primers (F1-GGGAGTTGGGGCAAATAATCC and R1-AAAACCTCCCACACCTCCC) were designed spanning the entire transgene integration cassette (4,889 base pairs) using Primer3web (version 4.1.0). New England BioLabs Q5 high-fidelity DNA polymerase was used following the manufacturer’s protocol for a 25-μl reaction volume using 5-10ng of gDNA. Thermal cycling was performed in a Bio-Rad C1000 thermal cycler using the following conditions: initial denaturing at 98°C for 30 s followed by 35 cycles of denaturing (98°C for 10 s), annealing (66°C for 30 s), and extension (72°C for 2:00 min) and a final extension at 72°C for 2 min. Annealing temperature was based on NEB Tm Calculator recommendation using default primer concentration of 500 nM. The resulting amplicon of the PCR was 4,889 nucleotides (nt). All PCRs included a positive control and a PCR reagent negative control. A total of 10 to 15μl of a PCR product was used for gel visualization.

### High content screening of transgenic larvae

Transgene inheritance frequency in F_1_ progeny was measured using a semi-automated, high content live-imaging approach (42). F_0_ animals were spawned and outcrossed with wildtype gametes. Embryos were imaged at gastrulation (24 - 26 hpf) by deciliation with 0.25 M NaCl and plating in a 96 well imaging plate (P96-1.5H-N, Cellvis). Images were taken in transmitted light and CFP fluorescence (445 nm wavelength, 60μm pinhole, exposure 100ms, illumination power 500mW) using the ImageXpress Spinning Disk Confocal HT.ai (Molecular Devices, San Jose, CA, USA) with a Nikon CFI Plan Apo Lambda 10X air objective. Machine learning-based automated segmentation of embryo z-stack max projection images (IN Carta v2.2, Molecular Devices) was used to count embryos and quantify embryo fluorescence as previously described(42). Embryos were binned as “integrated” or “nonintegrated” based on a maximum fluorescence of greater than or less than 235, and outliers with max fluorescence >1000 were excluded from analysis. To reduce false-positives, we removed “edge case” embryos with maximum fluorescence intensity between 230 - 275 from our analysis ensuring we only assessed correctly binned embryos. The rate of germline transmission (n_integrated_ / n_total_) was calculated for each F_0_ parent. Representative max projections were edited in Fiji by changing look up tables to cyan hot, adjusting max and min values, and adding a scale bar (49).

### Confocal Image Acquisition

Sea urchin larvae were mounted on a glass slide using clay feet for spacing between cover slip and slide. Larvae were mounted live in FSW with approximately 1:4 volumes of 1M MgCl_2_. Images were taken on a Leica TCS SP8 laser scanning confocal (APO 20x/0.70 CS dry objective) using a HyD detector at a 448 nm wavelength. Imaging parameters were set to a pinhole size of 2 AU, line average 4, and gain 30.0%. For larger larvae, the tilescan function of the Leica Application Suite X 3.5.7.23225 was used, merging together images with a 20% overlap. Juvenile sea urchins (0.5-1mm) were imaged live in FSW with approximately 1:4 volumes of 1M MgCl_2_ in Cellvis 96 well glass bottom plate (#1.5 high performance cover glass 0.17±0.005mm). Prior to adding the MgCl_2_, the animals were incubated in 3mL of FSW containing 0.3μl of stock concentration (5mg/mL) CellMask plasma membrane orange (Invitrogen) for 15 minutes. Images were taken on a ImageXpress Spinning Disk Confocal HT.ai (Nikon CFI Plan Apo Lambda 10X air objective) using CFP (445 nm wavelength, 60μm pinhole, exposure 100ms, illumination power 500mW and TRITC (555 nm, exposure 50ms, illumination power 500mW) filters. Images were edited in Fiji by projecting z-stacks with max projection, look up tables changed to cyan hot, max and min values adjusted, and scale bar added (49, 50).

### Fluorescence screening and statistical analysis

Larval screening was done using three to four different mate pairs with 50-100 injected embryos per mate pair totaling 200 embryos for each timepoint (day 2-16) and experimental group (+ *Minos* transposase, - *Minos* transposase).

Welch’s t-test were performed between timepoint (day) and experimental group to determine significant differences. Transposon testing at the juvenile phase was performed using at least 5 different mate pairs of injected embryos for each experimental group. Each experimental group (*Minos, Tol2, piggyBac*, controls) had a total of at least 400 juveniles screened for fluorescence. Kruskal-Wallis rank sum test followed by a pairwise Wilcoxon rank sum test was performed to determine statistical significance between groups. Integration efficiency (n_fluorescent_/ n_total_) was calculated using a weighted average to account for sample size differences between mate pairs.

## Acknowledgments and funding sources

The authors would like to thank Alexia Armendariz-Baca, Rachel Metry and Lorelei O’Brien for assistance with animal husbandry. This work was supported by NIH OD 034075 to AH.

## Competing interests

The authors declare no competing interest.

## Data, Materials, and Software Availability

Plasmids used to generate transgenic animals in this study have been submitted to Addgene.

## Notes

### Competing Interest Statement

The authors have declared no competing interest.

