## Supplemental for "Stable germline transgenesis using the *Minos* Tc1/*mariner* element in the sea urchin, *Lytechinus pictus*"

### Supplemental Figures

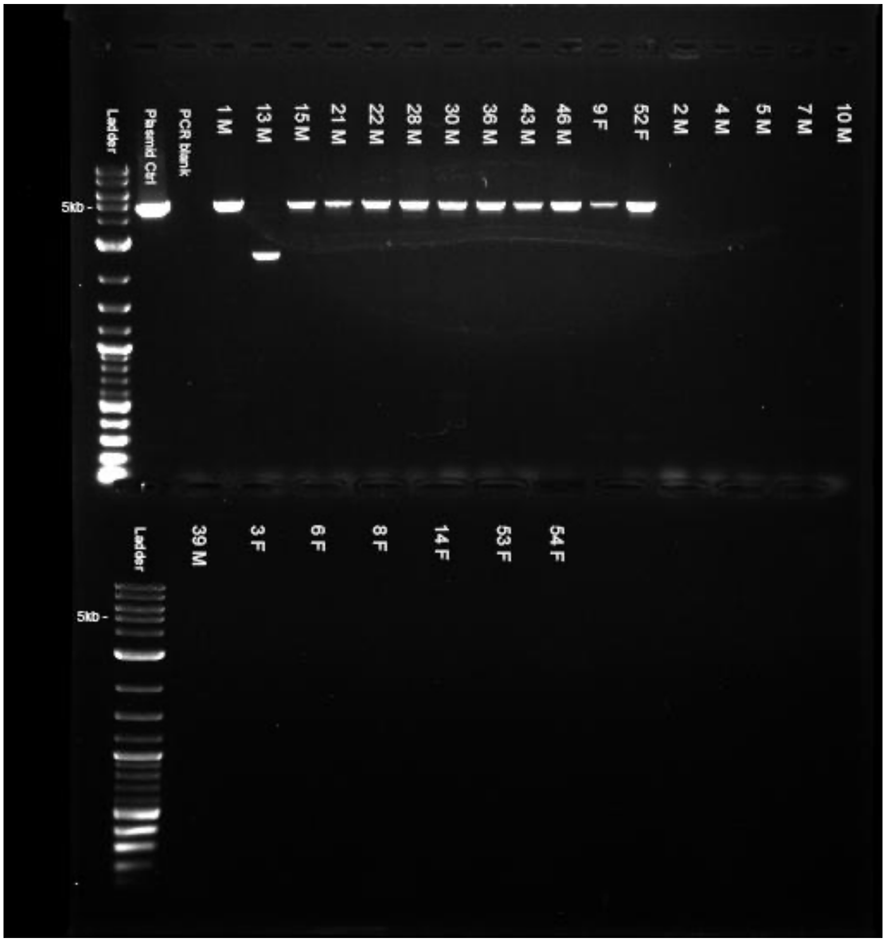


**Supplemental Figure 1. Representative PCR gel from DNA extracted from $F_1$ embryos.** $F_0$ adults were spawned and outcrossed with a wildtype male or female to determine germline integration. DNA extracted from $F_1$ embryos at ~24 hours post fertilization. The number corresponds to the animal ID. M = male and F = female.

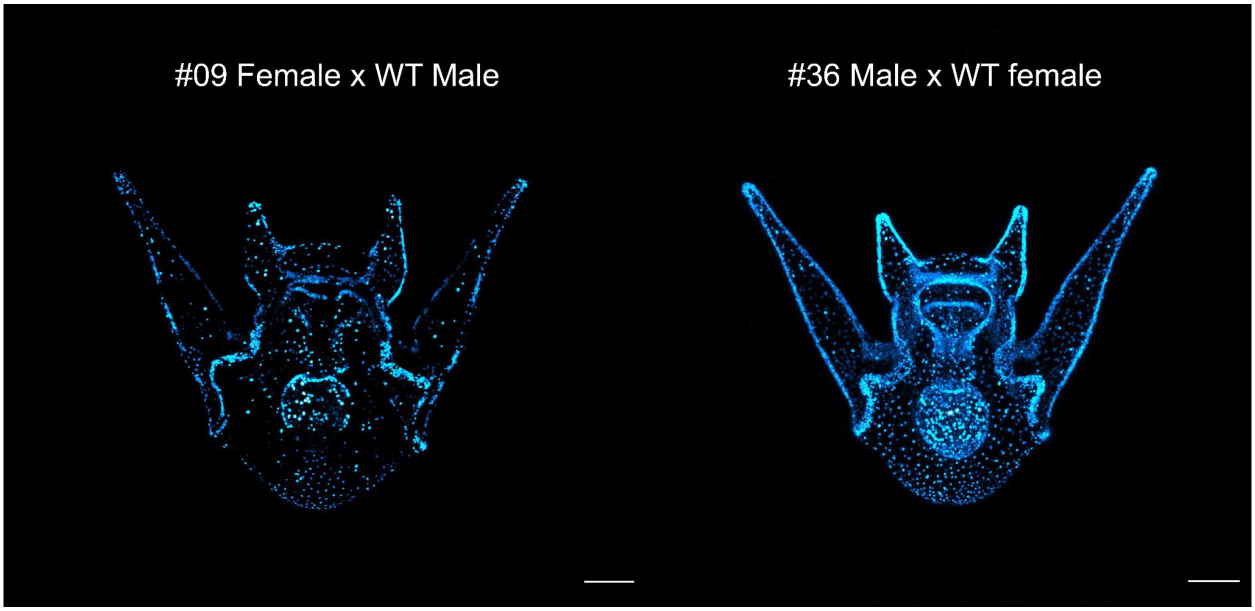


**Supplemental Figure 2. Difference in variegation of transgene expression in $F_1$ offspring.** The two females with germline integration in this study exhibited greater variegation of transgene expression compared to the males with germline integration. $F_0$ adults were spawned and outcrossed with a wildtype male or female. Larvae imaged at 7-8 days post fertilization. Scale bars = 100μm.

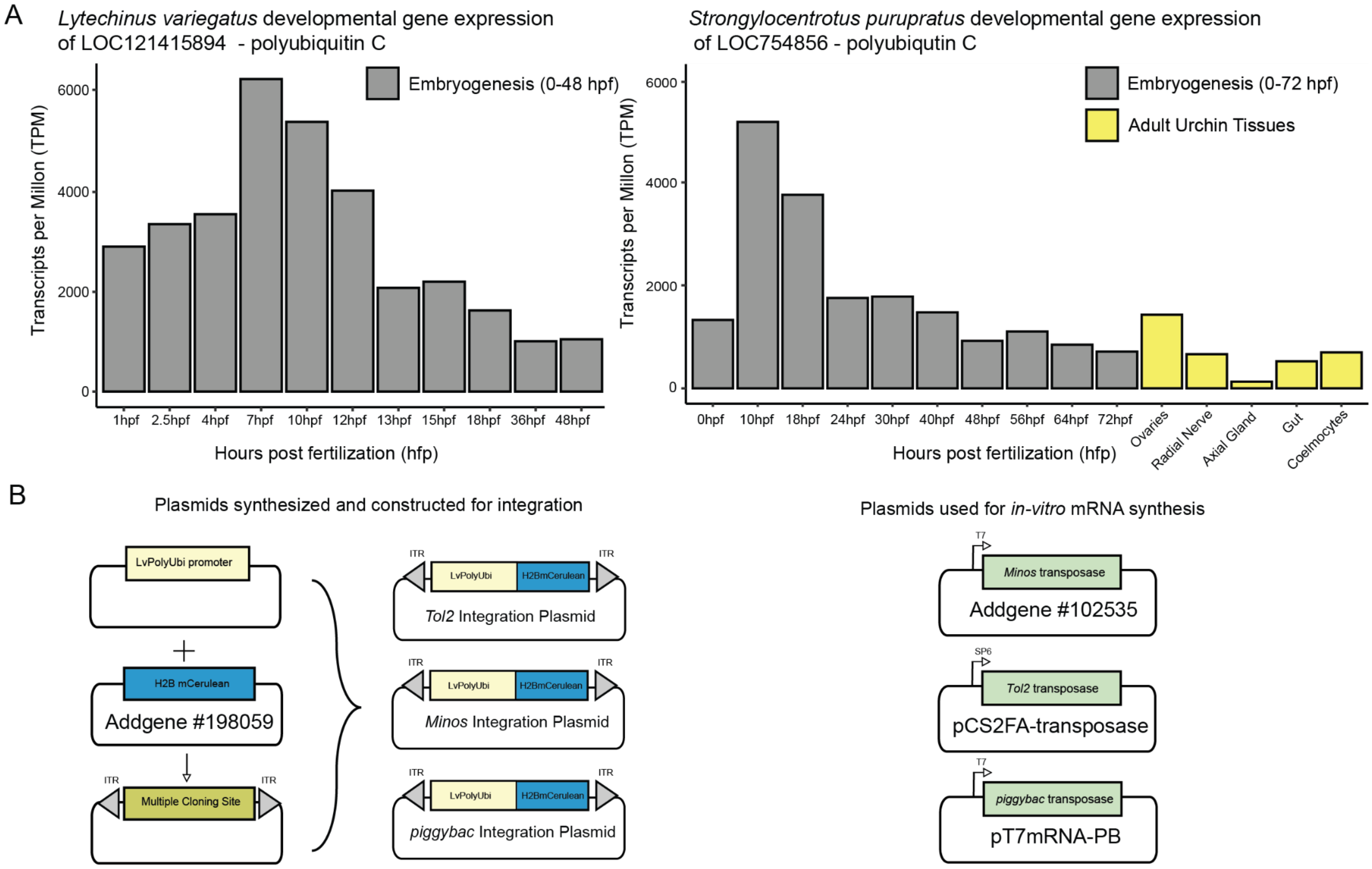


**Supplemental Figure 3. Promoter discovery and plasmid design for transposon-mediated integration testing.** (A) Gene expression pattern of polyubiquitin *L.variegatus* and *S.purpuratus* over early development, (B) Plasmids used and generated for integration.
